# Assessing the population genetic structure and demographic history of *Anopheles gambiae* and *An. arabiensis* at island and mainland sites in Uganda: Implications for testing novel malaria vector control approaches

**DOI:** 10.1101/2025.05.19.654785

**Authors:** Rita Mwima, Tin-Yu J. Hui, Edward Lukyamuzi, Marilou Bodde, Alex Makunin, Krystal Birungi, Martin Lukindu, Ann Nanteza, Dennis Muhanguzi, Mara Lawniczak, Austin Burt, Jonathan K. Kayondo

## Abstract

Despite substantial investments in malaria control, the disease remains a major burden in sub-Saharan Africa, particularly Uganda. Novel tools such as gene drive systems are being developed to suppress malaria vector populations, but their deployment requires detailed knowledge of mosquito population genetics.

We assessed the genetic structure, diversity, and demographic history of *Anopheles gambiae* and *Anopheles arabiensis* from six sites in Uganda: three islands in Lake Victoria and three mainland sites. A total of 2918 *Anopheles gambiae* and 173 *Anopheles arabiensis* were genotyped using targeted amplicon sequencing of 62 loci across coding and non-coding regions of the genome.

Population structure analyses revealed clear separation between the two species but little differentiation within each species across sites. Pairwise *F*ST values among *An. gambiae* populations were low (0.00054–0.028) but often significant, with mainland populations showing higher connectivity and island populations exhibiting greater isolation. *Anopheles arabiensis* mainland populations showed no statistically significant differentiation, suggesting panmixia. Principal Component Analysis and Bayesian clustering similarly distinguished species-level structure but no obvious substructure within sites.

Mainland *An. gambiae* populations displayed higher nucleotide diversity than island populations, while *An. arabiensis* showed the lowest diversity overall. Tajima’s D values were negative across sites, consistent with recent population expansions. Effective population size estimates indicated small populations at the islands (146 to 249) compared to large mainland populations (4,054 to 8,190).

These findings demonstrate strong genetic differentiation between *Anopheles gambiae* and *Anopheles arabiensis*, and subtle but meaningful structure between island and mainland *Anopheles gambiae* populations. The reduced diversity and small effective population sizes at island sites suggest stronger genetic drift and limited gene flow, in contrast to the highly connected mainland populations.

For malaria control, this contrast has direct implications. High connectivity among mainland populations may facilitate the spread of insecticide resistance alleles, while island populations, with their relative isolation and smaller sizes, may serve as suitable sites for contained field trials of gene drive strategies. This study highlights how geographic and ecological factors shape mosquito population structure and provides critical evidence for the design and monitoring of genetic-based vector control interventions.

## Background

Despite enormous efforts in malaria control, the disease remains a global burden, especially in the Sub-Saharan Africa. This is because of the exceptionally adaptive dynamics of *Anopheles* mosquitoes together with other factors that include insecticide resistance, drug resistance, limited access to healthcare, and environmental and socio-economic challenges [1,2].

In 2024, the World Health Organisation (WHO) reported about 200 million cases and 600,000 deaths [3]. Recent years have shown a persistently high malaria burden with case and mortality numbers remaining largely stable from 2021 to 2023, underscoring the health challenge the disease poses, despite the control efforts in place [4,5]. Sub-Saharan Africa carries the greatest burden globally, accounts for 94% of all malaria cases and 96% of all deaths [3]. Uganda, in particular, experiences a heavy burden due to malaria, ranking third in the number of malaria cases worldwide in 2023 and contributing 5% of the global burden [3].

The fight against malaria is increasingly challenged by the evolving resistance of parasites to antimalarial drugs and of mosquitoes to various insecticides, coupled with the partial efficacy of current control methods [3,6–9]. To achieve malaria elimination and eradication will thus require new products and interventions to be used in tandem with the already existing tools [10]. One promising strategy is the mosquito gene drive technology, a genetic engineering approach that biases the inheritance of a specific gene or trait, causing it to spread rapidly through a population over generations, capable of spreading into an entire population from low initial frequencies [11,12] to either suppressing mosquito populations by reducing their fertility or modifying them so they are unable to transmit the malaria parasite [13].

Significant advancements have been made so far in target gene identification, gene construct development, and genome sequencing of the major malaria mosquito populations [14–16].

Despite the proof-of-concept mosquito transformation studies done in laboratories and the idententification of candidate gene drive systems [17–19], there is need to advance for further field trials such that product feasibility and efficacy can be evaluated. Furthermore, detailed population genetic assessments must be done systematically before deploying any proposed trial or intervention [20]. Information on population genetic structure and gene flow within a vector species provides a basis for modeling the impact of trial intervention and in trial design and helps select suitable locations for field trials, given that, Ideally, these locations should have limited gene flow with surrounding populations [21,22].

In Uganda, the geographic and genetic isolation of lacustrine *An. gambiae* island populations from mainland renders them suitable candidate sites for initial testing of mosquito gene drive systems [15]. Conversely, regions with high gene flow are significant for gene drive strategies because they provide high connectivity among populations and enhance the potential for rapid and extensive spread of gene drive elements, which is essential for effective malaria vector control on a broader scale [23–25].

Knowledge of A*n. gambiae* and *An. arabiensis* genetic diversity, structure, and population sizes at island and mainland sites will provide critical insights into the genetic differences and dynamics at the target field sites, thereby aiding in the design of interventions and monitoring their efficacy [21,26,27]. Several studies have examined the structure of *An. gambiae* at the island and mainland sites in Uganda [28–30], but there are no studies that compare the population structure of both *An. gambiae* and *An. arabiensis*, despite their known sympatric occurrence in many regions [31,32]. These two species are closely related members of the *Anopheles gambiae* complex, sharing overlapping habitats and breeding sites, which creates opportunities for gene flow and introgression that could impact important traits such as insecticide resistance and vector competence [21,33]. Therefore, assessing their population structure and potential gene flow jointly, is critical to understanding their ecological interactions and for optimizing vector control strategies. *Anopheles arabiensis* from the lacustrine islands is often excluded from further analysis due to their limited numbers in the mosquito collections done to date [26,27,34], and yet anecdotal observations suggest that the islands on Lake Victoria may have an increasing presence of *An. arabiensis* alongside *An. gambiae*, suggesting a shift in species composition and complexity of malaria vector species in these areas (personal communication). The explanation for this could be This could be, microclimatic conditions that result in increased density of a given species in areas where their numbers were previously very low [35–37].

This study employed population genetics approaches like meansures of genetic diversity, differentiation and gene flow estimates to characterize the population structure of *An. gambiae* and *An. arabiensis* across selected island and mainland sites in Uganda. This analysis utilised amplicon data generated from an amplicon panel specifically designed for species identity.

## Methods

### Mosquito sampling

Mosquito collections were conducted from six sites including three mainland sites (Kayonjo, Katuuso, Kibbuye) and three island (Bugiri, Kiimi, Kansambwe) sites (Figure 1). These sites were chosen based on several criteria: (1) high malaria endemicity, (2) a high density of the target mosquito species (*An. gambiae* complex), (3) a distance from urban areas with a mix of permanent and temporary housing, (4) a moderate population size (between 500-1000 residents), (5) minimal ongoing malaria vector control activities, and (6) a cooperative and welcoming local community. Kibbuye and Katuuso villages are situated in Mukono district and Kayonjo village in Kayunga district (Figure 1), which are each approximately 50km North West of the capital city, Kampala. The lacustrine island sites are situated within Lake Victoria in Uganda and are connected by ship and ferry service to the Ugandan mainland. Figure 1B shows how far each site is from another.

**Figure 1.**
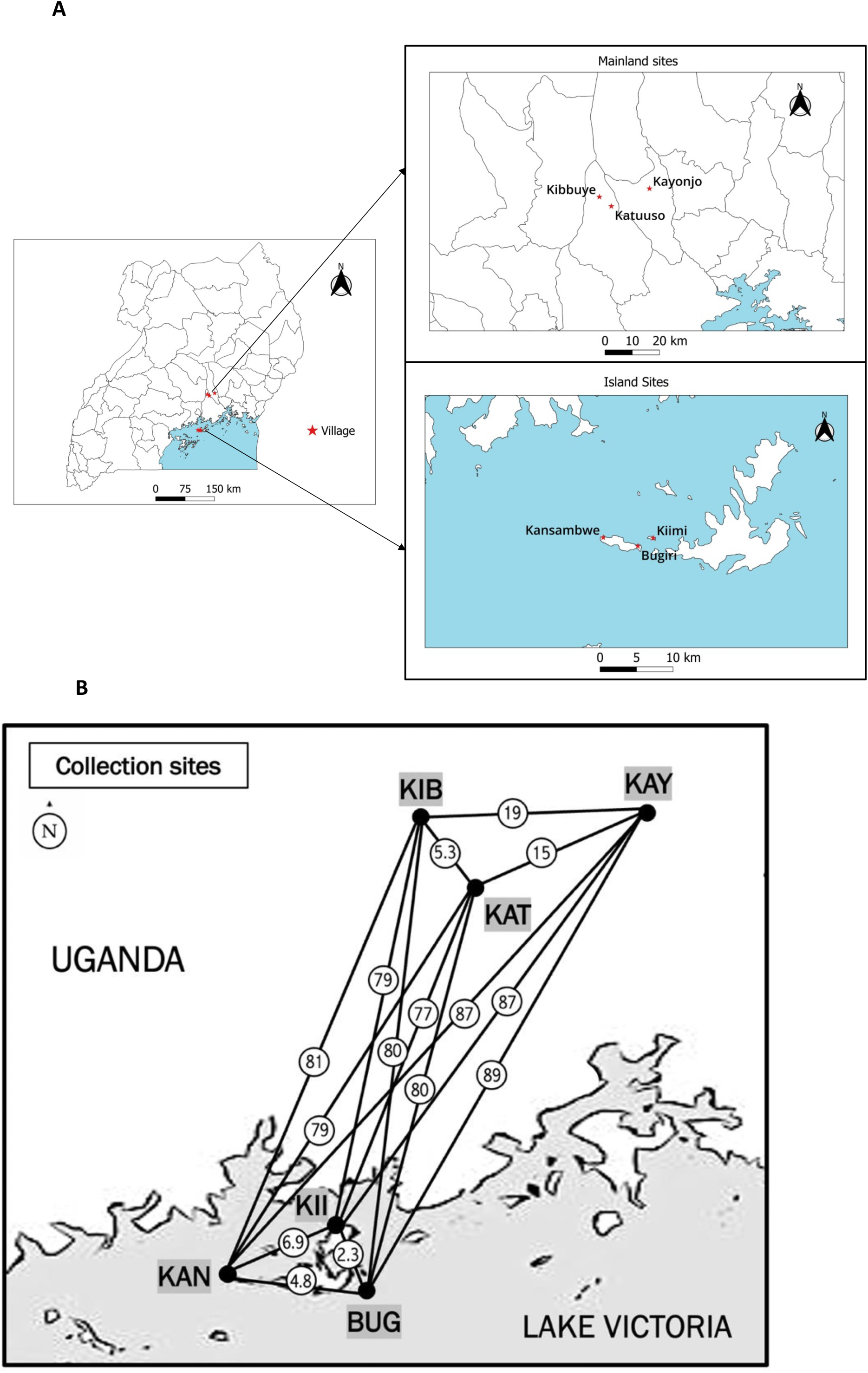
A: Map showing the location of the 3 mainland and 3 island study sites in Uganda, **B:** Distances (in kilometers and circled) between sites. KAY= Kayonjo, KIB = Kibbuye, KAT = Katuuso, KII = Kiimi, KAN = Kansambwe, BUG = Bugiri.

Collections done on the islands and mainlands were part of routine mosquito collections done by the Department of Entomology at the Uganda Virus Research Institute. On the islands, collections were done from April 2013 to April 2016, and on the mainland from January 2016 to December 2018. Adult mosquitoes were sampled indoors and outdoors using human landing catches (HLC), pyrethroid spray catches (PSC), indoor and outdoor aspirators, and catch basin traps (CBT). All mosquitoes were morphologically identified using the *Anopheles* morphological identification keys [38] and then stored in 80% ethanol for subsequent molecular analysis. For this study we kept only *An. gambiae* and *An. arabiensis* mosquitoes, although some *An. funestus*, *An. maculipalpis*, and *An. coustani* were also collected (but excluded from this analysis).

### Genomic DNA extraction

A total of 3515 whole mosquito samples, each stored in a 96-well plate of 80% ethanol, were shipped to the Wellcome Sanger Institute, United Kingdom, for DNA extraction and amplicon sequencing [16,39]. In brief, DNA extraction was completed by removing the ethanol from each specimen, adding 100 ul of lysis buffer C to each, and incubating plates at 56 ^0^C for overnight, allowing preservation of specimens for further examination [16]. DNA from each mosquito was then subjected to a single Polymerase Chain Reaction (PCR) reaction containing a 64 primer pair plex [16]. Each reaction then went into a second PCR to add indexing primers. Eight plates of mosquitoes were pooled for a single MiSeq library [16].

Using the ANOpheles SPecies and Plasmodium (ANOSPP) panel of 64 phylogenetically informative and highly variable “amplicon loci” of which 62 target the Anopheles nuclear genome and the other 2 targeting the Plasmodium mitochondria (note: these amplicon loci are of ∼160 base pairs or “sites” long in downstream analyses), the generated sequence data enabled species identification, detection of Plasmodium, hybrid or contaminated samples, and identification of cryptic species [16]. This panel consists of 62 nuclear loci distributed across the *Anopheles* genome, comprising 17 coding (exonic), 22 intronic, and 23 intergenic (non-coding) regions [16]. (For all other details regarding the sequencing techniques, and the functions of the 62 amplicon loci, please refer to the original publication [16,39]).

A Variational Autoencoder (VAE), which is a machine learning approach that identifies patterns in high-dimensional data by compressing it into a lower-dimensional representation [40] was used to distinguish between closely related species [39].

### Bioinformatics processing

The raw reads from the generated sequence data for the confirmed *An. gambiae* and *An*. *arabiensis* were processed using AmpSeeker, an open-source computational pipeline which includes quality filtering, primer trimming, alignment to the reference genome file (VectorBase-59_AgambiaePEST_Genome.fasta), and variant calling (e.g with BWA, bcftools). [40]. AmpSeeker is a Snakemake-based workflow designed for Illumina amplicon sequencing data [41]. Using this pipeline, samples were filtered based on sequencing depth, heterozygosity and principal component outlier analysis to remove low-quality or anomalous samples, before downstream analyses like population structure and gene flow were performed [41].

The resulting raw variant call format (VCF) file contains variant calls across all samples and comprises a total of 12,412 sites (inclusive of insertions and deletions (indels), primer regions, fixed and polymorphic sites). Subsequent subsetting was performed on the VCF file based on the metadata information, which provided details about the mosquito samples, including individual sample IDs, species, collection location, collection period, and season among others. In the filtering process, indels were removed, resulting in 9,890 remaining sites. These sites were distributed across the genome as follows: X (n = 952), 2L (n = 1881), 2R (n = 2815), 3L (n = 1615), 3R (n = 2627). The minor allele frequency (maf) spectrum between 0 and 2% is shown in Figure 2, and the number of alleles per locus (from fixed to quadriallelic) is in Figure 3. Our downstream analyses were based on these 9,890 sites with potential further filtering, such as according to maf and missing values (Using only biallelic SNPs).

**Figure 2:**
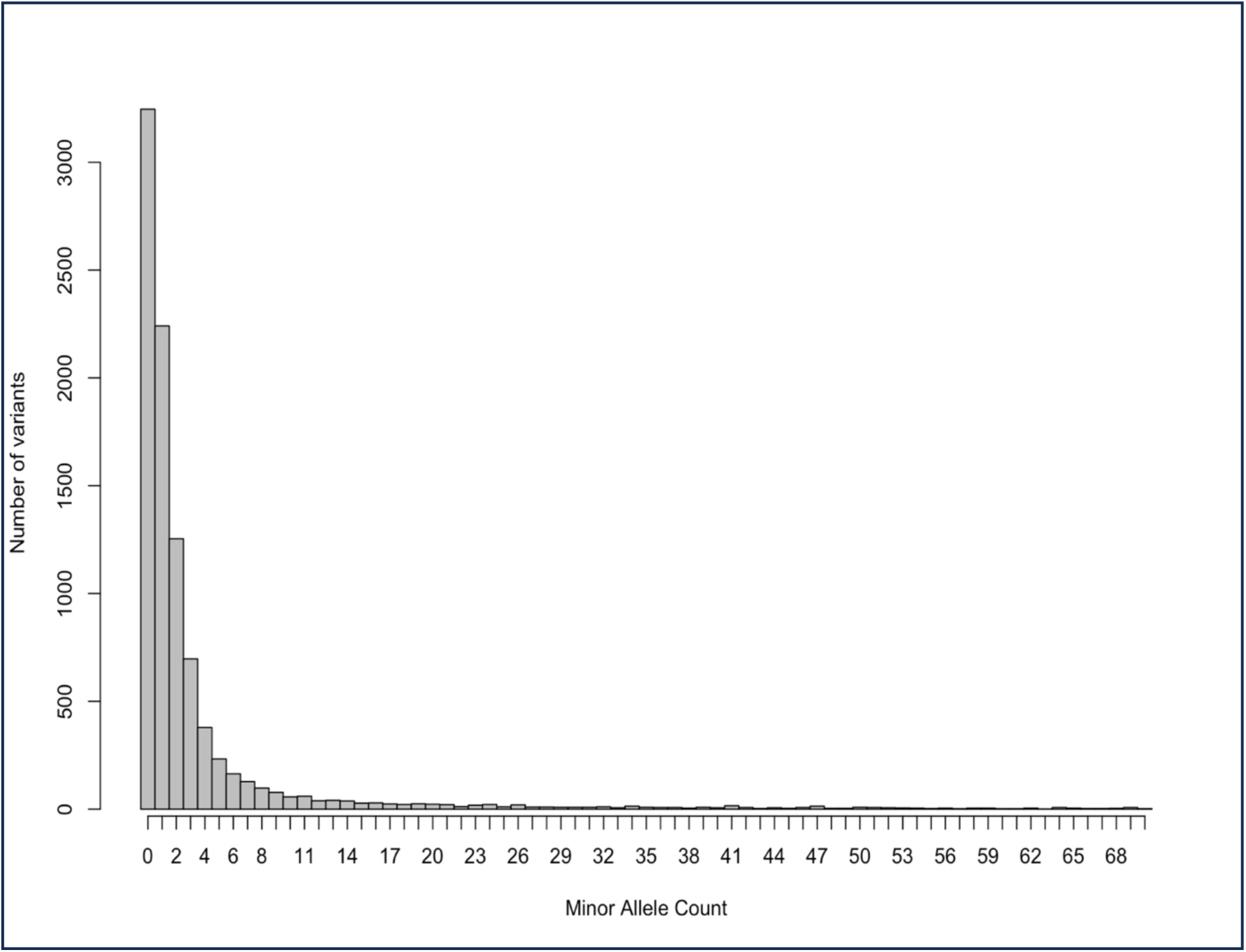
Minor allele frequency (MAF) spectrum between 0 and 2% (for the whole dataset).

**Figure 3:**
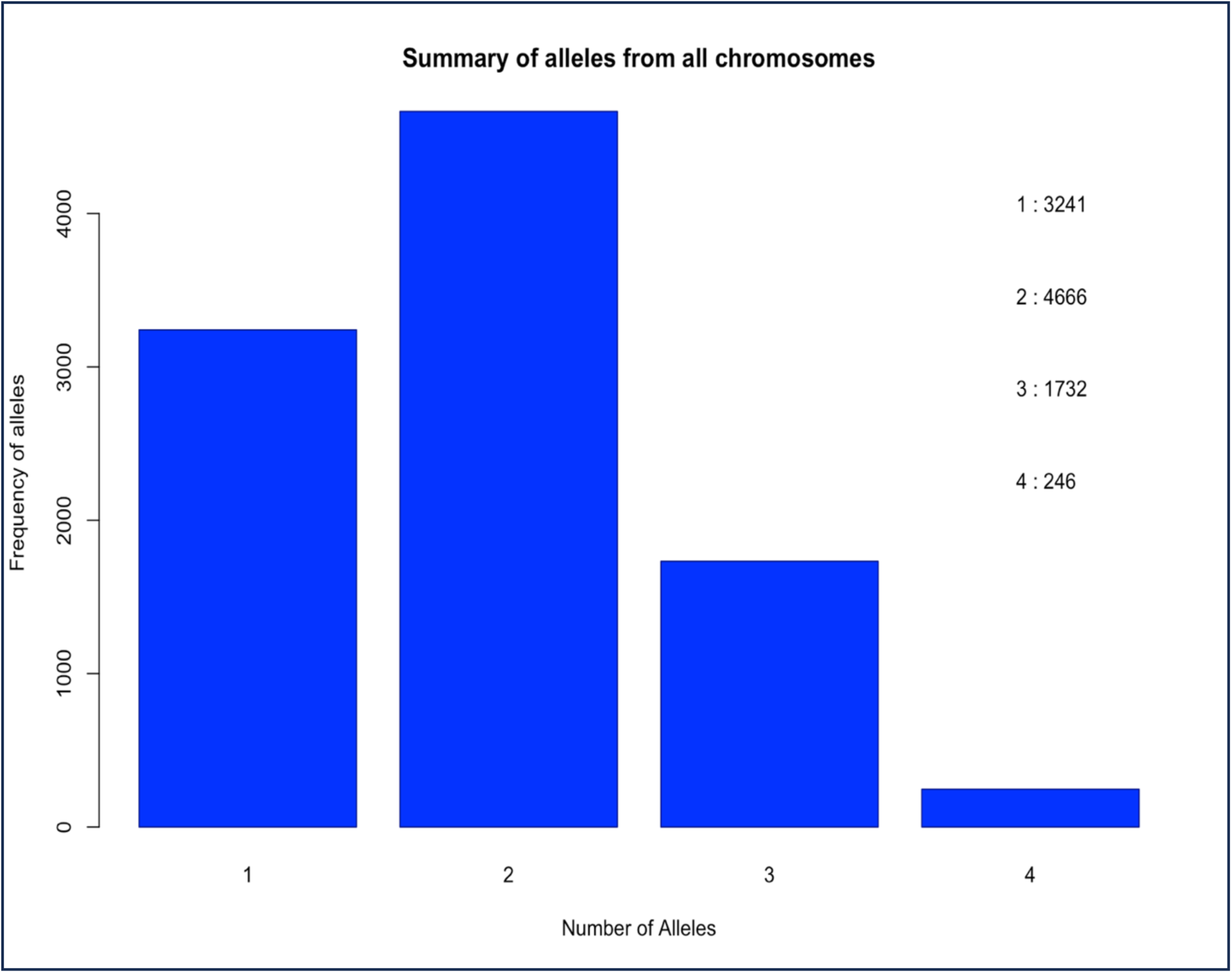
Summary of the number of alleles per site from all chromosomes. 9890 sites are included. The 5 sites with 0 alleles are missing values in all our *An. gambiae* and *An. arabiensis* samples and were excluded from the graph.

## Population genomic analysis

### Population structure

The following methods were used to describe population structure: (1) *F_ST_* between and within pairs of populations using version 1.3.23 of the Popkin package in R [42,43], (2) Principal Components Analysis (PCA) using version 3.6.2 of the prcomp package in R [42,43], and (3) Bayesian clustering analysis with STRUCTURE v2.3.4 [44]. Average pairwise Hudson’s *F_ST_* was calculated using the fst_hudson_pairwise function of the Popkinsuppl package [45], after which standard errors were computed using the leave-one-out approach that is a specific form of the jackknife resampling method [46–48]. Z-scores were then computed by dividing the estimated FST by its standard error. These Z-scores were used to assess the significance (P-values) of the *F_ST_* values. The P-values are derived from the Z_scores by comparing them against the standard normal distribution, which allows for determining the probability of observing such *F_ST_* values under the null hypothesis of no differentiation between populations [48]. All the *An. gambiae* collected from the same location were considered as a population, while all 173 *An. arabiensis* were grouped as the seventh population. The *An. arabiensis* samples were grouped as a single population in the analysis because their sizes per site were too small to conduct statistically robust population-level analyses individually. Grouping all *An. arabiensis* samples increased the analytical power to detect genetic patterns and structure across the mainland populations, providing a more reliable overview of their genetic differentiation.

For pairwise *F_ST_*, we used biallelic sites with 10% missing data from within the 3L chromosome with maf >=1%, while for PCA, we first considered all chromosomes and thereafter only the 3L arm, because, unlike other autosomes, it does not carry chromosomal inversions that suppress recombination and could confound analyses of population structure. To investigate the role of geographic distance in shaping genetic differentiation, we ran a Mantel test to test for isolation by distance (IBD) between populations by combining Euclidean geographic distances (calculated from geographic coordinates) and genetic distances based on 9999 permutations using GenAlEx 6.51b2 [49,50]. For *An. arabiensis* populations, the same *F_ST_* and Mantel tests were also conducted using the 3L chromosome arm biallelic SNPs but only for the three mainland sites. The islands were left out because of their small sample sizes (Table 1). PCA was performed in R v4.3.1 with the prcomp() function [51,52], first for biallelic SNPs from all chromosomes and then for the 3L chromosome arm.

**Table 1.** The number of confirmed *An. gambiae* and *An. arabiensis* samples across six collection sites.

|  | Bugiri † | Kiimi † | Kansambwe † | Kayonjo | Katuuso | Kibbuye | Total |
| --- | --- | --- | --- | --- | --- | --- | --- |
| *An. gambiae* | 28 | 258 | 294 | 1178 | 450 | 710 | 2918 |
| *An. arabiensis* | 2 | 0 | 3 | 51 | 39 | 78 | 173 |
| Total | 30 | 258 | 297 | 1229 | 489 | 788 | 3091 |

† Denotes island sites.

We looked at doubletons, defined as loci with exactly two copies of the minor allele in a biallelic site. A contingency table of observed counts of heterozygous doubleton pairs across populations was constructed, and the expected counts were calculated based on sample size proportions to assess population structure. Populations were grouped into three categories, *arabiensis* (*An. arabiensis*)*, mainland_gam* (mainland *An. gambiae*), and *Island_gam* (island *An. gambiae*), and a chi-square test was conducted to compare observed and expected counts to detect significant population structure.

To find the optimal number of clusters (K) in STRUCTURE (using model with no admixture), we conducted 20 independent runs for each K (from 2-8), using a burn-in value of 100,000 iterations followed by 100,000 repetitions [44]. Then K was determined using the Delta K method of Evanno et al. [53], following which CLUMPAK was used to construct a graphical representation of the genetic structure of the 3091 (both *An. gambiae* and *An. arabiensis*) mosquito samples [54]. In this analysis, we randomly chose one variable site per amplicon locus from the 2R, 3R, and 3L chromosome arms to minimise genetic linkage, with the constraint that maf≥1%.

### Genetic diversity and Ne

To assess the genetic diversity per population, we calculated the overall nucleotide diversity (p) as well as individually for the 6 *An. gambiae* and 3 mainland *An. arabiensis* populations for each chromosome arm [55] using VCF files without missing data. Deviation from the standard neutral model was tested using Tajima’s D, which was performed for each site and chromosome to examine population expansion [56]. Tajima’s D and nucleotide diversity were computed using the Pegas package in R [57]. Estimates of contemporary Ne were attained using the linkage disequilibrium (LD) based method LDNe [58] of NeEstimator v.2.01 [59]. We randomly chose one site per amplicon locus from chromosomes 3R and 3L with maf≥1% to avoid tight linkage. We similarly individually computed nucleotide diversity, Tajima’s D and contemporary Ne for individual mainland *An. arabiensis* populations.

## Results

### Genetic differentiation (F_ST_)

2918 samples were confirmed as *An. gambiae* and 173 as *An. arabiensis*. These individuals and their resulting amplicon data were our subjects for downstream population genetic analyses, and Table 1 summarises their distribution across the six collection sites. The average pairwise Hudson’s *F_ST_* is found in Table 2A. All pairwise comparisons produced low *F_ST_* values (0.00054-0.028), except between the “outgroup” *An. arabiensis* and any *An. gambiae* populations. Among *An. gambiae* mainland populations, pairwise *F_ST_* values were very low (0.00054-0.0016, indicating minimal genetic differentiation, and similarly, low *F_ST_* values were observed between Bugiri (island) and the other two islands (Kiimi and Kansambwe), suggesting significant gene flow. The highest *F_ST_* values occurred between Kansambwe (island) and all the mainland sites (Kayonjo, Katuuso and Kibbuye) (island vs mainland), and between Kiimi and Kansambwe (island vs island), implying restricted gene flow in these comparisons (Table 2A). The pairwise *F_ST_* values between Bugiri (island) and all the other sites, Kayonjo (mainland) and Katuuso (mainland), Kiimi (island) and Katuuso(mainland) and Kiimi (island) and Kibbuye (mainland) were low and non-significant (z < 1.96, P > 0.05). We then calculated pairwise *F_ST_* values among the mainland *An. arabiensis* populations (Table 2B), and low and statistically nonsignificant pairwise *F_ST_* values (-0.0014994 - 0.00785541) were mostly recorded. The *F_ST_* value between Katuuso (mainland) and Kibbuye (mainland) was negative, which is interpreted as no differentiation.

**Table 2:** Pairwise *F_ST_* statistics.

|  | BUGIRI † | KIIMI † | KANSAMBWE † | KAYONJO | KATUUSO | KIBBUYE | *An. arabiensis* |
| --- | --- | --- | --- | --- | --- | --- | --- |
| BUGIRI † | 0 | 0.6796089 | 0.5914296 | 1.762527 | 1.628254 | 1.47858 | 4.361302* |
| KIIMI † | 0.00287 | 0 | 4.631849* | 1.977574* | 1.943631 | 1.78073 | 4.360677* |
| KANSAMBWE † | 0.00272 | 0.02123 | 0 | 3.189523* | 3.041091* | 3.916723* | 4.041615* |
| KAYONJO | 0.00995 | 0.01199 | 0.02715 | 0 | 1.204753 | 1.991724* | 3.779667* |
| KATUUSO | 0.01201 | 0.01483 | 0.02841 | 0.00054 | 0 | 2.439927* | 3.826917* |
| KIBBUYE | 0.01033 | 0.01740 | 0.02433 | 0.00109 | 0.00163 | 0 | 3.737203* |
| *An. arabiensis* | 0.21710 | 0.20209 | 0.26279 | 0.22654 | 0.23330 | 0.20902 | 0 |

|  | KAYONJO | KATUUSO | KIBBUYE |
| --- | --- | --- | --- |
| KAYONJO | 0 | 1.817574 | 0.09362245 |
| KATUUSO | 0.00785541 | 0 | -1.070926 |
| KIBBUYE | 0.00018109 | -0.0014994 | 0 |

(A) Pairwise $F_{ST}$ from the 3L chromosome arm for the six *An. gambiae* populations, with the
173 *An. arabiensis* grouped as the seventh population. (B) Pairwise $F_{ST}$ among the three *An.*
*arabiensis* populations. For both tables, the lower left triangle in each table shows average
Hudson’s $F_{ST}$ values between each population pair, while the upper right triangle shows the z-
scores for each $F_{ST}$ value calculated via a block-jackknife method. The values with \* indicate significant $F_{ST}$ values ( $z > 1.96$ , $p < 0.05$ ). † Denotes island sites.

The pairwise *F_ST_* values for *An. gambiae* were plotted against geographic distances to investigate the strength of isolation by distance [50,60], and the correlation was not statistically significant (Figure 4, P = 0.052).

**Figure 4:**
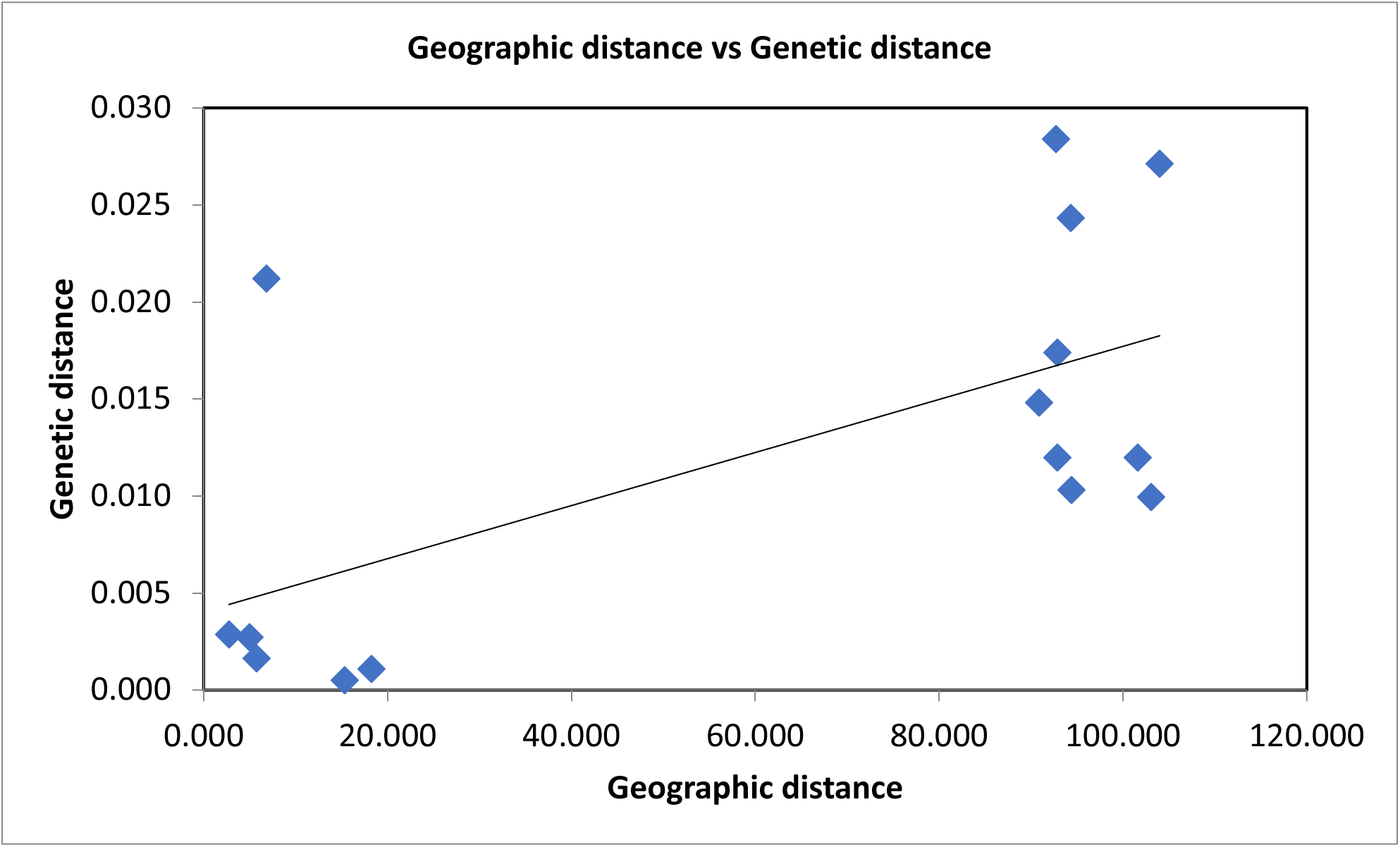
F*_S_*T versus geographical distance (km) for *An. gambiae*. FST values were extracted from Table 2. The figure shows a positive slope with no significant correlation between the genetic and geographic distances, P = 0.052

### PCA

PCA visualises how the individuals were genetically clustered within and between collection sites. Analysis using all biallelic sites of all chromosome arms of all *An. gambiae* and *An. arabiensis* samples showed clustering into two distinct groups in accordance with each species’ genetic difference (Figure 5A). The 3L chromosome arm from both *An. gambiae* and *An. arabiensis* also showed species-specific clustering (Figure 5B) and using chromosome 3L sites of only *An. gambiae*, no geographical clustering of individuals was observed (Figure 5C).

**Figure 5.**
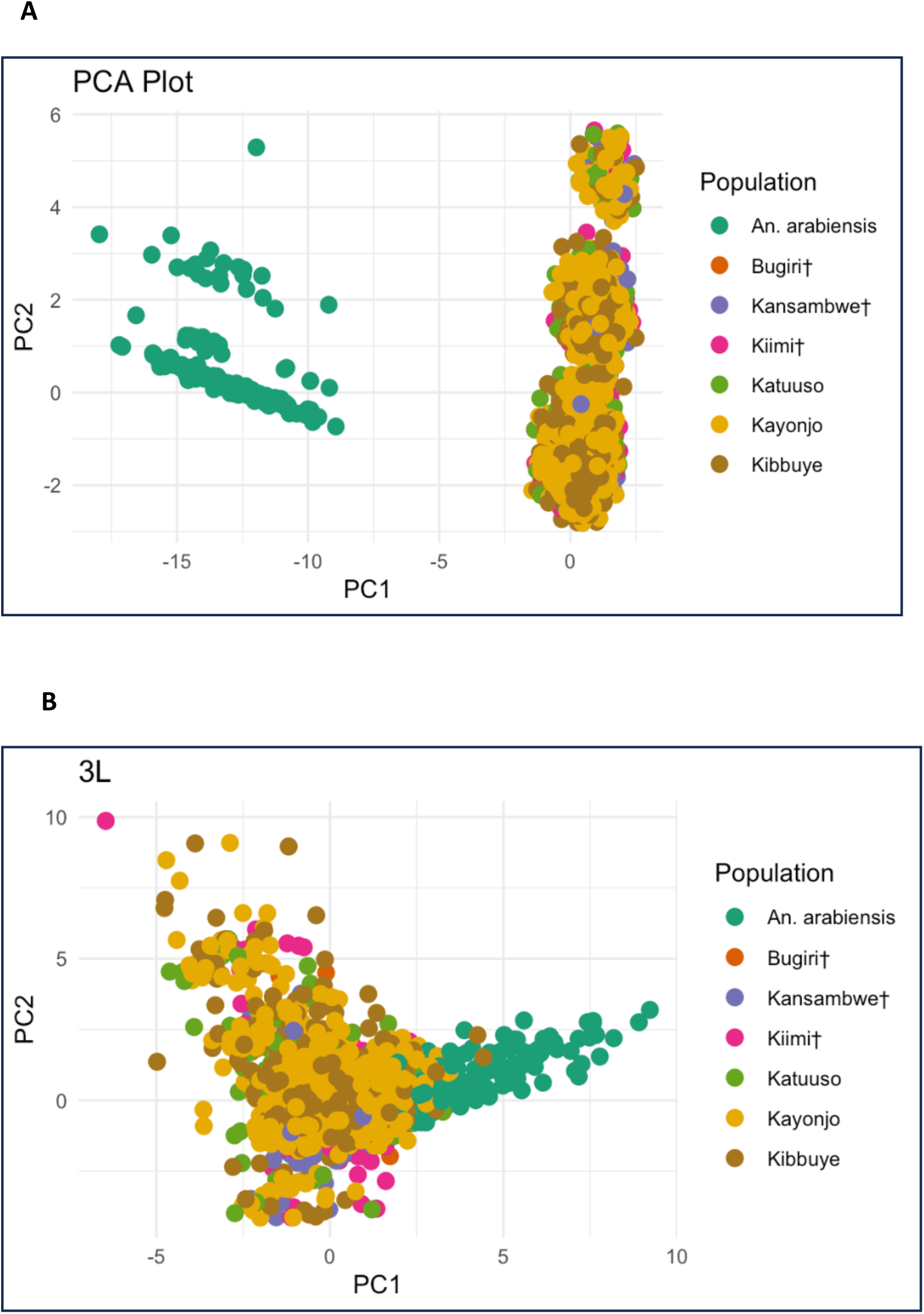
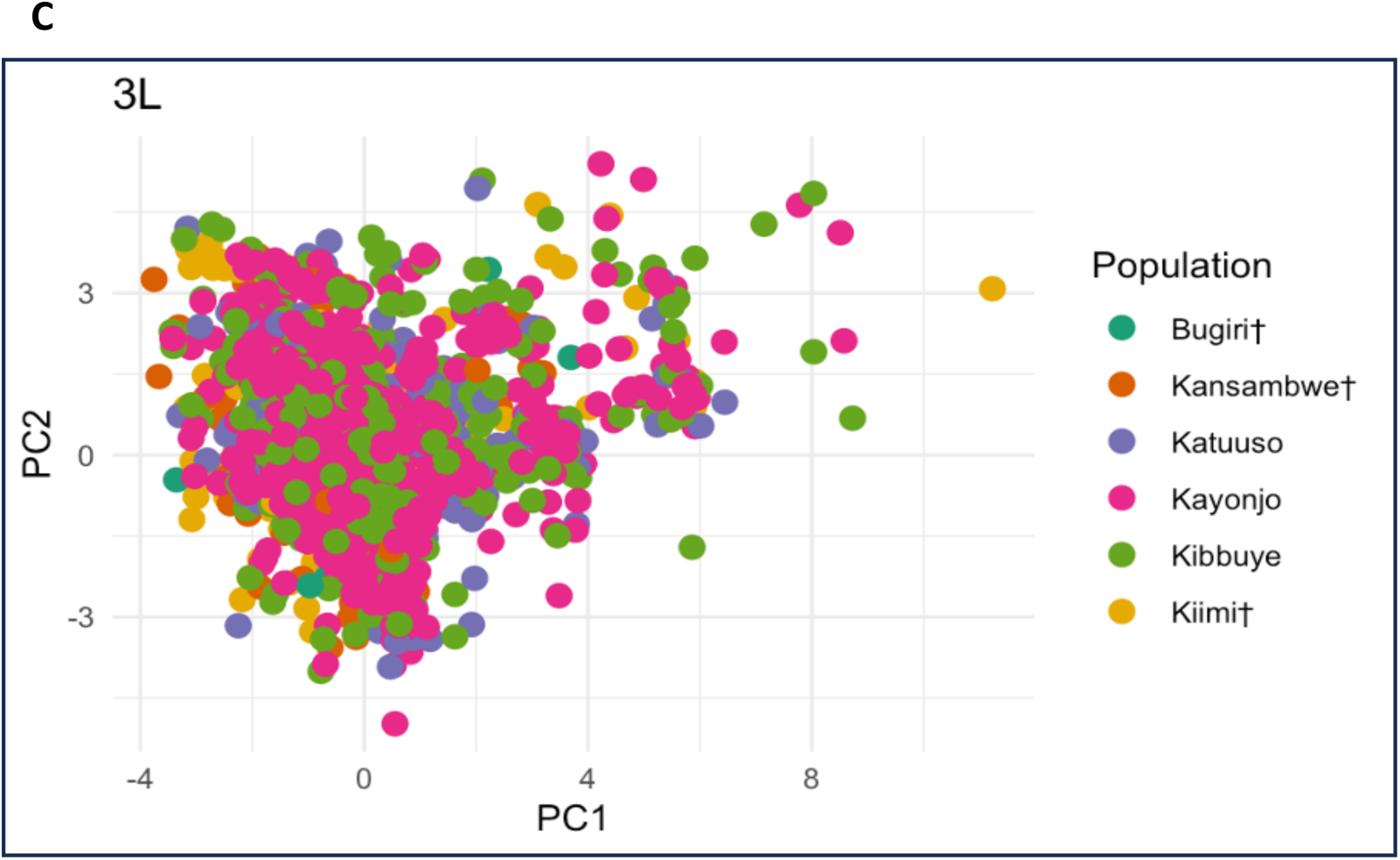
A: Principal component analysis of all chromosome arms of *An. gambiae* (island and mainland sites) and *An. arabiensis***, B:** The PCA was calculated from the 3L chromosome arm of *An. gambiae* and *An. arabiensis* individuals, and **C**: The PCA was calculated from the 3L chromosome arm of only *An. gambiae* individuals. † Denotes island sites.

### Allele sharing in doubleton variants

839 pairs of doubletons were discovered. We calculated the expected counts based on sample size proportions to account for the distribution of the observed heterozygote doubleton pairs under a null hypothesis assuming no population structure. Because of the limited resolution available to distinguish between individual populations in our dataset, we reduced the original 6+1 (Bugiri, Kiimi, Kansambwe, Kayonjo, Katuuso, Kibbuye, and *An. arabiensis*) population groups into three broader categories: *arabiensis*, *island_gam*, and *mainland_gam*. In this way, more robust statistical comparisons and interpretations of allele sharing across major population segments could be made. The reduction from 6+1 populations to three groups improved statistical power and provided insights into gene flow dynamics across the major geographic and species categories of interest.

There were significant deviations between observed and expected pairwise counts of doubleton pairs in the three groups: *arabiensis*, *island_gam*, and *mainland_gam*. The observed pairwise counts of heterozygotes within and between population groups highlight actual genetic similarities, while the expected pairwise counts reflect the hypothetical distribution of pairings in the absence of genetic structuring.

When we considered within-group genetic structuring (Table 3), the *mainland_gam* group showed a significantly higher within-group count of heterozygous pairs than expected (682 observed vs. 480 expected), which suggests a higher-than-anticipated allele sharing or pairing frequency within this group. The *island_gam* group showed a lower-than-expected level of allele sharing (2 observed vs. 29.54 expected), suggesting limited gene flow with other populations, while, the *arabiensis* group showed moderated within-group genetic isolation (15 observed versus 2.63 expected counts), suggesting that it is a relatively closed genetic pool with reduced gene flow to other populations.

**Table 3:** Allele sharing in doubleton variants.

A. Observed counts
|  | *arabiensis* | *island_gam* | *mainland_gam* |
| --- | --- | --- | --- |
| *arabiensis* | 15 | 4 | 102 |
| *island_gam* | NA | 2 | 34 |
| *mainland_gam* | NA | NA | 682 |

B. Expected counts
|  | *arabiensis* | *island_gam* | *mainland_gam* |
| --- | --- | --- | --- |
| *arabiensis* | 2.628186 | 17.62252 | 71.03699 |
| *island_gam* | NA | 29.54064 | 238.1587 |
| *mainland_gam* | NA | NA | 480.01296 |

Within and between group counts

When we considered between-group genetic structuring, the observed gene flow between *arabiensis* and *mainland_gam* groups (102 observed versus 71.04 expected counts) indicates occasional genetic exchange, which could be as a result of ancestral similarities. *island_gam* versus *mainland_gam* have significantly fewer pairs than expected (34 pairs versus an expectation of 238.16 pairs), suggesting restricted gene flow between the two geographic sites. The *arabiensis* versus *island_gam* groups had significantly fewer pairs than expected (4 pairs versus an expectation of 17.62252 pairs), suggesting restricted gene flow between the two groups.

### Bayesian clustering analysis

Based on the log-likelihood values and the DeltaK plot, Κ=3 was the optimal number of evolutionary clusters (Figure 6A). Therefore, the six *An. gambiae* populations and pooled *An. arabiensis* were substantially admixed at Κ = 3 (Κ = 3 best explains the genetic variance present between all the sites). Upon close inspection of the results from the Bayesian cluster analysis(Figure 6B), three significant genetic clusters were identified: (1) pooled *An. arabiensis* individuals, (2) island *An. gambiae* individuals, (3) mainland *An. gambiae* individuals. *An. arabiensis* were assigned to the orange population with the highest probabilities, separating them from all *An. gambiae* mosquitoes. The island *An. gambiae* were mostly assigned to the blue but with slightly more uncertainly. Much of the variation was recorded between populations (*An. gambiae* island and mainland individuals and *An. arabiensis* individuals) instead of among individuals within a site. These results from STRUCTURE (Figure 6B) were also largely consistent with those from PCA (Figure 5A-C) with clear separation at an interspecific level and no clear separation at the intraspecific level.

**Figure 6:**
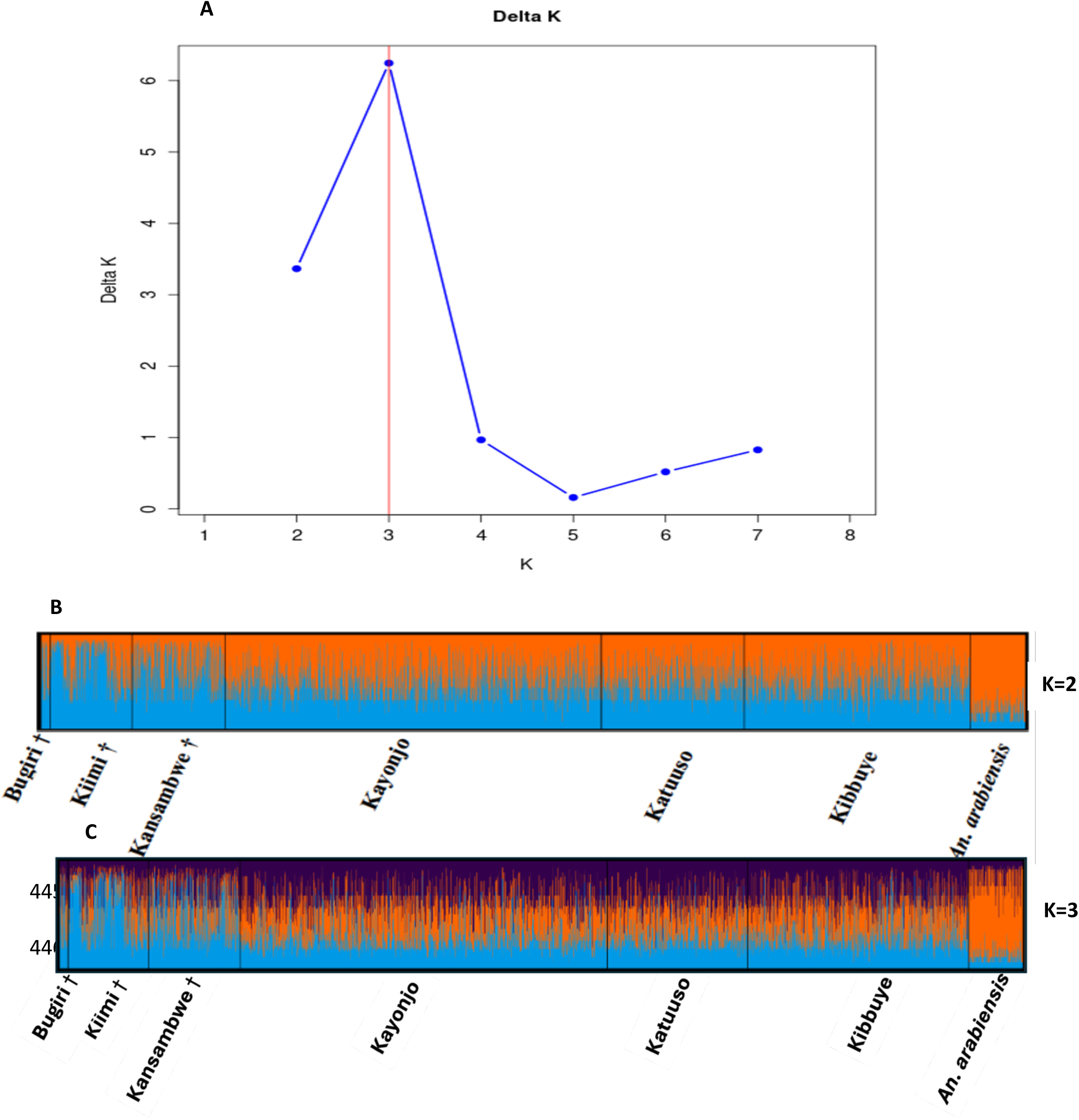
Outputs from Bayesian clustering analysis for the 2918 *An. gambiae* and 173 *An. arabiensis*. **(A)** The optimal number of clusters (*K=3*) based on the Evanno et al [61] method showing deltaΚ **(B)** Bar plot representing K=2 of 3091 mosquitoes, each represented by a thin vertical bar colored in proportion to their estimated ancestry within each cluster (**C**) Bar plot representing K=3 of 3091 mosquitoes. A vertical black line bounds each site and the labels with † for islands while those without are for mainland samples.

### Genetic diversity and neutrality tests

Nucleotide diversity (π) and Tajima’s D were calculated for each *An. gambiae* population (SI Table S1), then further for each chromosome arm (Figure 7A). The overall nucleotide diversity for *An. gambiae* was 0.0115 (range from 0.0107 to 0.01169, SI Table S1) and that of *An. arabiensis* was 0.00859 (ranging from 0.00829 to 0.00859; SI Table 2). Generally, higher diversity values were reported for mainland *An. gambiae* populations (0.01143 to 0.01169; SI Table 1) compared to those for island sites (0.01071 to 0.01103; SI Table 1). On the other hand, the diversity values for *An. arabiensis* mainland populations were lower compared to the mainland *An. gambiae* populations. The pattern of nucleotide diversity compared between chromosomes for different sites (Figure 7B) was consistent across all sites and chromosome arms (mainland sites’ p > island sites’ p). At each site, mean chromosomal nucleotide diversity was highest on the 3R chromosome and lowest on the 2R arm. However, pooled *An. arabiensis* (from all locations) had an overall low diversity compared to the different *An. gambiae* populations, with the lowest diversity in the 2R chromosome arm (P = 0.031, Figure 7B). Overall, the pooled island and mainland diversity falls between the values for each sample site across chromosome arms (Figure 7B).

**Figure 7:**
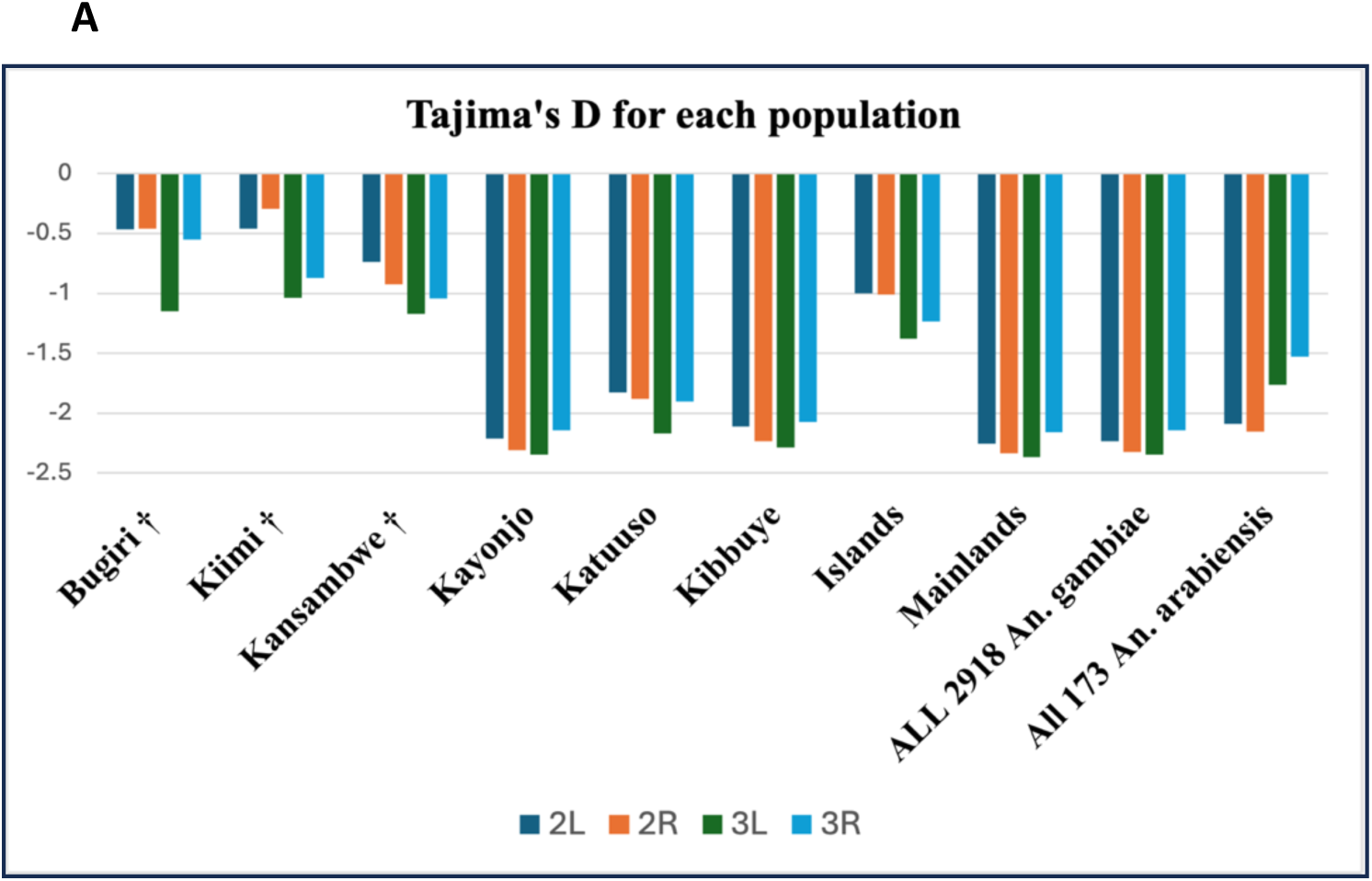
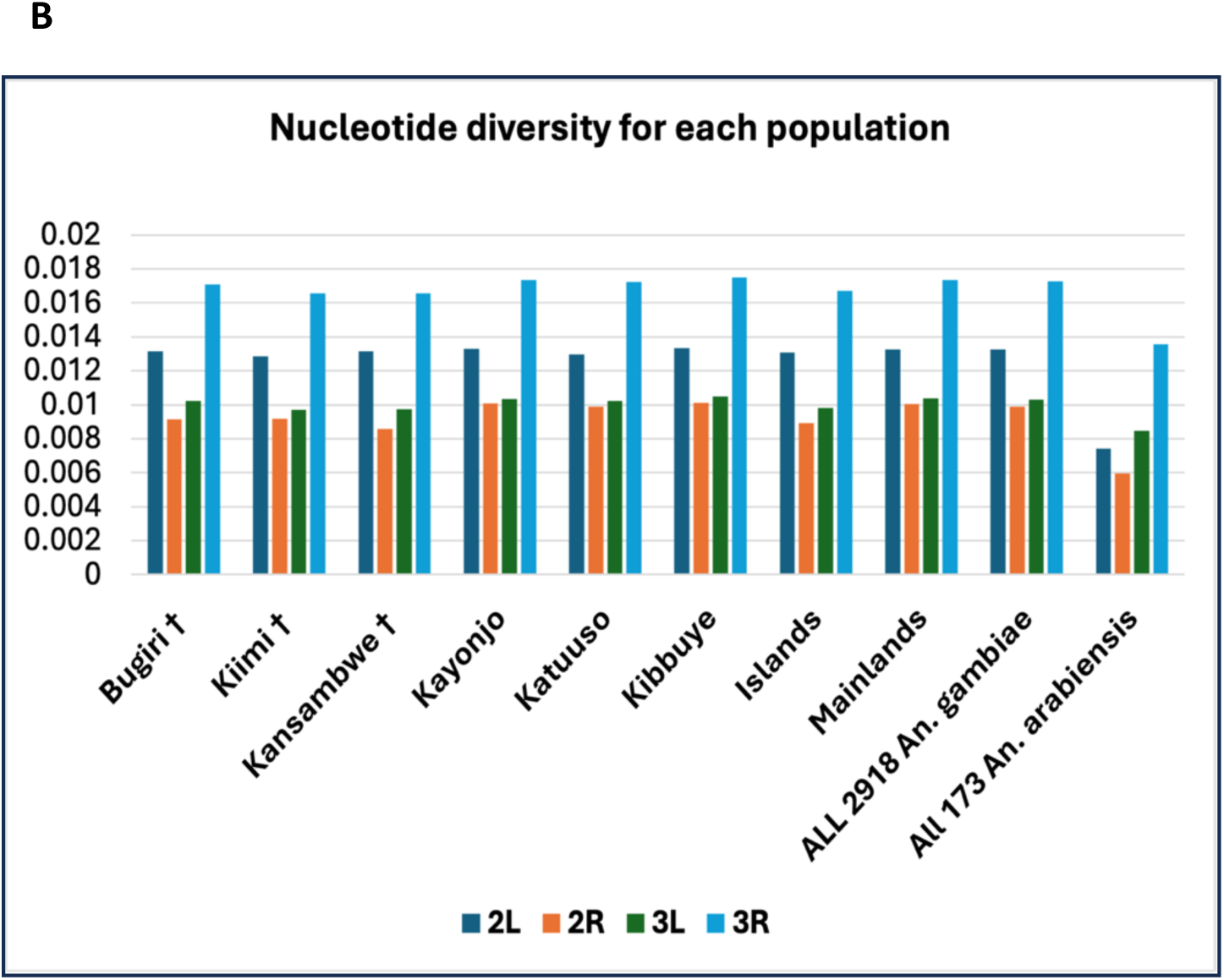
**(A)**Tajima’s D for the six *An. gambiae* populations per chromosome arm. In addition, we calculated the same statistic for the following subsets: pooled *An. gambiae* from the island and the mainland sites, then all *An. gambiae*, and all *An. arabiensis* (**B)** Nucleotide diversity (π) for the six *An. gambiae* populations per chromosome arm. † Denotes an island site.

Mean Tajima’s D was negative for all sites, an indication of an excess of rare alleles and thus a deviation from the neutral model of a well-mixed population of constant size. However, unlike the *An. gambiae* mainland populations, Tajima’s D values for *An. gambiae* island sites and all *An. arabiensis* populations were not statistically significant from from zero (SI Table S1, S2). When tested for each chromosome and site, all Tajima’s D values were negative. The same was done for the *An. arabiensis* mainland populations (SI Table S2). The Tajima’s D value of the pooled *An. arabiensis* chromosome 2R arm was significantly more negative than other arms of the same site (P = 0.031), and the 2R arm of the individual and pooled populations.

### Contemporary effective population size (*Ne*)

Generally, for *An. gambiae* populations, smaller Ne values were observed for the island sites (between 146 and 249), compared to mainland sites (between 4054 and 8190) (Table 4A). The point estimate for Bugiri was inconclusive but came with a very small lower bound. For the *An. arabiensis* populations, we only estimated Ne for the mainland sites, and their estimates were comparable to the mainland *An. gambiae* populations. (Table 4B).

**Table 4:**
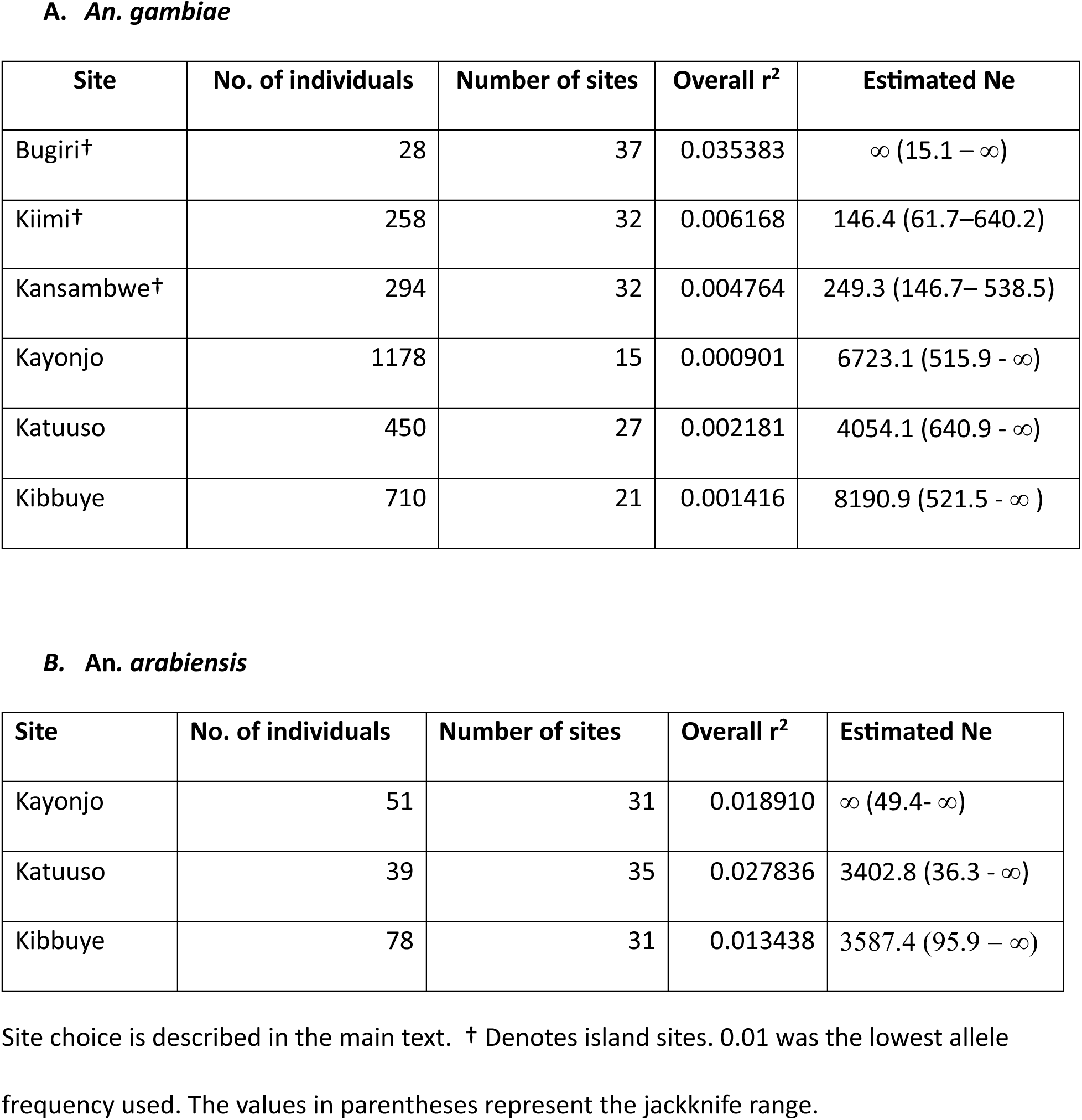
Contemporary Ne estimates.

## Discussion

### Polymorphic sites and population structure

From the 62 amplicon loci, 9890 sites were mapped onto the reference genome, many of which were fixed sites (Figure 3) or weakly polymorphic due to the conserved nature of the primer-binding regions (Figure 2). Despite this limitation, the data revealed useful insights into the population structure of *An. gambiae* and *An. arabiensis* in Uganda. Overall, genetic differentiation among the *An. gambiae* populations was generally low regardless of the sites (*F_ST_* < 0.05), though statistically significant in some cases, suggesting subtle structure with ongoing connectivity.

The high gene flow observed between Bugiri (island) and other sites could be influenced by several factors. One potential cause is human-mediated transport, as frequent boat traffic and ferries between Bugiri and other sites can inadvertently facilitate mosquito dispersal [62–64]. Additionally, prevailing seasonal wind and weather patterns might assist mosquito flight over water, enhancing connectivity [65,66]. These mechanisms together contribute to the low genetic differentiation reported for Bugiri compared to other island populations. However, the low *F_ST_* value observed for Bugiri could also be influenced by its relatively small sample size, which can increase sampling variance and reduce the power to detect subtle genetic differentiation [67,68].

In contrast, *An. arabiensis* mainland populations exhibited very low and often negative *F_ST_* values, consistent with unrestricted gene flow [46,69,70]. The negative *F_ST_* values (between Katuuso and Kibbuye) likely reflects greater within-site genetic diversity than between-sites genetic diversity, indicative of stable and large Ne [71,72].

One notable feature of our results is the presence of low but statistically significant *F_ST_* values among many population pairs, indicating that while the genetic differentiation is subtle, structure nonetheless exists. *F_ST_* values for island *An. gambiae* species were however higher compared to mainland sites, reflecting restricted gene flow across water barriers, smaller population sizes, and potential local adaptation comparable to what was reported in previous studies [26,27,34]. This could be justified by the large mainland population, given that *F_ST_* is approximately 1⁄1 + 4𝑁𝑚, where 𝑁 is the effective population size and 𝑚 the migration rate [72]. Thus, using our estimated large mainland populations for N, the low *F_ST_* values between sites equate to substantial numbers of migrants per generation, reinforcing connectivity among sites [68,72]. Given that these *F_ST_* values are significantly greater than zero demonstrates that the populations are not panmictic or identical but exhibit genuine population structure [73,74]. Recognizing this subtle but meaningful structure is crucial when considering the design and spread dynamics of gene drive or insecticide resistance alleles.

Looking at this from a malaria vector control perspective, high gene flow is a double edged sword. On one hand, it promotes the spread of gene drive constructs throughout connected mosquito populations, potentially enabling rapid spread of desired genetic traits [75]. Conversely, it complicates containment, given that, genetic modifications could unintentionally spread beyond target areas, raising ecological and regulatory concerns. Moreover, convergent evolution may promote the rise of resistance alleles that impair gene drive function, necessitating careful monitoring [76].

It is thus important to distinguish between whether low differentiation results are from ongoing gene flow, convergent adaptation, or both, and understanding these dynamics allows for better-informed strategies to optimize gene drive deployment, balancing efficacy with biosafety in malaria vector control.

Low point estimates of pairwise fst yielded high Z_scores, suggesting the presence of subtle yet statistically significant population structure despite low apparent differentiation. This occurrence can largely be attributed to the statistical power afforded by a large sample size, which can detect even minor genetic differences across sites [77,78]. Between Kiimi and Katuuso, the low pairwise fst value and non-significant P-value could be because of temporal sampling variability, which could result in capturing different allele frequencies due to temporal fluctuations [79–81]. These comparisons of the relative amounts of gene flow taking place between sites, aid in predicting the trajectory of the alleles introduced by gene drive into the wild-type population [21].

The Mantel test results (P=0.052) suggested only weak evidence of isolation by distance [82], suggesting that geographic distance partly influences gene flow, which could thus be factored into spatially targeted vector control interventions [68]. Therefore, according to the IBD hypothesis, there may be a meaningful association between the genetic and geographical distances among populations. Therefore, tailored intervention deployment considering spatial genetic structure can optimise control effectiveness.

PCA distinguished species but did not reveal clear geographical structure within *An. gambiae*. These findings align with doubleton analysis (Table 3), which highlighted relative isolation of island sites and stronger connectivity at mainland sites.

The moderate within-group isolation of the *arabiensis* group (Table 3) indicates more frequent genetic interactions within this group than expected, potentially due to localized genetic structures, ecological factors, or other selective pressures driving allele frequencies. This could reflect a localized population structure which may arise from ecological factors like microhabitat variation, behavioural differences or localized selective pressures, that can restrict gene flow at finer spatial scales despite the species’ wide distribution.

The counts between *arabiensis* and *mainland_gam* groups indicate a degree of gene flow, implying occasional genetic exchange, which could have direct implications for vector control, especially given the differential patterns of gene flow and inbreeding detected across these sites. This pattern of observed versus expected doubleton pairs sides with the broader view of population structure in malaria vectors, where poor gene flow across ecological boundaries frequently results in discrete genetic clusters [83], and has implications on both gene drive strategies and management of insecticide resistance.

STRUCTURE analysis identified three major clusters (Figure 6) corresponding to species and geography, consistent with *F_ST_* and PCA results. STRUCTURE results reflect what is often observed when there is low genetic differentiation [84]. STRUCTURE output for the *An. gambiae* and *An. arabiensis* species at each site showed sharing of ancestry between each individual at differing proportions (with differing consistency between all island, mainland *An. gambiae* and *An. arabiensis* individuals). The three genetic clusters corresponded to first species (*An. arabiensis*) then to geographical (mainland and island *An. gambiae*) differences, with relatively more similarity between island compared to mainland *An. gambiae* individuals (Figure 6B). These results are comparable to the *F_ST_* and PCA results that show less divergence between island and mainland *An. gambiae* individuals and more diversity (clear separation) between *An. gambiae* and *An. arabiensis* individuals [55,56].

Despite using very few polymorphic sites, STRUCTURE was able to outline the three clusters with visible differences. Therefore, using more sites could produce clearer and more distinct clustering. And because pairwise *F_ST_* is sensitive to allele frequency differences between sites and works best when populations are distinct and genetically isolated, it may not always capture subtle population differentiation, while STRUCTURE, on the other hand, assigns individuals to clusters based on their multilocus genotypes, and is thus more sensitive to subtle population differentiation [44,73,85,86].

### Genetic diversity and neutrality tests

Negative Tajima’s D values (SI Tables S1&S2), especially at mainland sites, suggested an excess of low frequency alleles, potentially reflecting recent population expansion or purifying selection [87–89]. These results were in line with previous findings [83]. Given that *An*. *gambiae* thrives in human dorminated environments, its population and range will thus expand most in areas where human population density is high and growing, whether on island or mainland sites [90].

The observed differences in Tajima’s D and nucleotide diversity between island and mainland *An. gambiae* populations provide further evidence that these are distinct populations rather than a single panmictic unit. Mainland populations exhibited significantly higher nucleotide diversity and more negative Tajima’s D values, consistent with recent population expansion and larger effective population sizes [72,91]. In contrast, island populations showed marginally lower nucleotide diversity and Tajima’s D values not significantly different from zero (SI Table S1, S2), suggesting small but stable effective population sizes or mutation-drift equilibrium, with no population bottleneck or expansion, an occurrence consistent with the neutral mutation hypothesis [92,93]. These genetic differences, together with the subtle but significant population structure detected by *F_ST_* and clustering analyses, support the conclusion that island and mainland *An. gambiae* populations are genetically differentiated and should be considered as separate population units for vector control considerations [34,64].

These estimates were within the range of values reported in previous studies which utilised four loci (white, tox, G6pd, xdh) across several African locations [94] and immune-related (LRIM1, CTL4, CTLMA2, APL2) and housekeeping genes [95], and were consistent with the fact that populations at island sites display marginally lower nucleotide diversity compared to the mainland [96,97]. The explanation for this could be inbreeding, or higher degree of isolation with small neighborhood size [96,98–100].

On the 2R chromosome arm, the pooled *An. arabiensis* population has lower nucleotide diversity than individual or pooled *An. gambiae* populations, which could be because the 2R arm has many segregating inversions, some of which are polymorphic in *An. gambiae* and fixed in *An. arabiensis,* which inversions can limit recombination in certain regions, leading to reduced genetic diversity over time [101,102]. This phenomenon is especially pronounced in species like *An. arabiensis*, where these inversions (2La, 2Rb and 3Ra) could be fixed, and further decrease genetic variation along the 2R chromosome relative to *An. gambiae* populations [103,104].

Additionally, the frequencies for inversions segregating in both species could be/are different for the two species [102], suggesting a decline in population size and/or balancing selection, a feature of the maintenance of segregating 2Rb, c, d, u and j inversions [90,105].

### Contemporary effective population size

Ne estimates were smaller for island *An. gambiae* than at mainlands sites, reflecting isolation and stronger drift, results comparable to those reported in the Kayondo et al. and Wiltshire et al. studies [28–30]. The low Ne estimates at island sites could be because of their small neighbourhood size and are suggestive of higher levels of genetic differentiation (presented as pairwise F_ST_ estimates) shown in Table 4A [59].

Mosquito populations at these smaller, more isolated sites are more vulnerable to genetic drift, but may also be more amenable to localized control interventions.

The infinity values of point estimates such as those recorded for Bugiri, was almost certainly due to insufficient sample size compared to the magnitude of genetic drift. The infinite jackknife upper bound at the mainland sites suggests that based on the LD observed, it’s statistically feasible that Ne could be extremely large, although the exact size cannot be pinpointed with certainty [106–108].

Larger Ne in the mainland *An. gambiae* populations suggest higher resilience to vector control interventions and more likely to sustain the spread of resistance alleles [109–113]. This implies slower genetic drift which allows for sustained genetic variation that enhances adaptation to environmental changes, and because dry season surviving populations can rebound and maintain genetic diversity across seasons, this further complicates efforts to control malaria vector populations [114–118].

### Implications for malaria vector control

These findings have important implications for: (1) insecticide resistance: high connectivity among mosquitoes at mainland sites facilitate rapid spread of resistance alleles, where as island isolation may slow spread and allow for localised resistance management, (2) Gene drive strategies: the reduced Ne and relative isolation of mosquitoes at island sites lower the release threshold for gene drives, increasing the likelihood of successful establishment [21]. Relative isolation and smaller Ne values favours islands as contained experimental sites for gene drive trials, allowing more manageable monitoring and containment [75]. Conversely, large, well-connected mainland sites would require larger releases and careful management to ensure effective drive propagation, and (3) Vector elimination efforts: The weak but detectable structure among *An. gambiae* populations suggests that control efforts need to account for ongoing gene flow between sites. Local elimination on islands may be feasible, but reinvasion from the mainland remains a threat.

The ease of access to the islands (which allows easy monitoring), higher genetic differentiation, genetic structure, smaller effective population sizes compared to mainland sites, and their small geographical size, makes islands are promising sites for field trials to test the effectiveness of mosquito gene drive systems.

And as mentioned at the start, *An. arabiensis* individuals are often excluded from further analysis due to their limited numbers in the mosquito collections done to date [26,27,34], which could be because of the sampling methods used. Therefore, studies that target the exophilic and zoophagic characteristics of this species should be carried out to give a better picture of its density, specifically at the islands which are the potential sites for field trials to test mosquito gene drive systems. This will help us to confirm the proportion of the non-target malaria vector species at the island sites.

As much as we would like to provide a comprehensive perspective of the overall spatial and temporal variation of the two species in Uganda, the latter cannot be investigated here because of the different sampling times in the island and mainland sampling. We welcome further studies on other biological factors, such as the mosquitoes’ seasonal dynamics and dry season persistence mechanisms [119,120], as these may contribute to the observed population structure.

Future works should include a dataset using whole genome sequenced data, exploring temporal sampling, further investigations into the specific contribution of the suggested factors we have used, as well as the influence of other suggested biotic (e.g., passive transport) and abiotic factors (e.g., seasonality and chromosome inversions) that may contribute to the observed differentiation. These will add a valuable body of information required for evaluating these Islands as potential release sites for mosquito gene drive systems.

## Conclusion

Despite limited polymorphism in the loci studied, this work highlights subtle but meaningful population structure between *An. gambiae* and *An. arabiensis* populations. The contrast between small, isolated island sites and large, connected mainland sites has direct consequences for the design of insecticide resistance management strategies, feasibility of gene drive interventions, and broader malaria vector control approaches.

## Acknowledgements

We are grateful to Cindy Sewon, a former undergraduate student at Imperial College, London for her valuable contributions, John B. Connolly, Silke Fuchs and Samantha O’Loughlin for their support and helpful comments towards this work, and the field team that did the mosquito sample collection.

## Conflict of interest

The authors declare no conflicts of interest. The funding bodies had do direct role in the design of the study nor in the collection, analysis, interpretation of data or in the writing of the manuscript

## Author contributions

JK conceived the research topic, hosted lab and field work. ML hosted the targeted amplicon sequencing. MB and AM participated in amplicon sequencing. EL supported in bioinformatics processing. RM and TH analysed the data. RM wrote the manscript with inputs from all other authors. All authors read and approved the final manuscript.

## Funding

This work was supported by the Bill & Melinda Gates Foundation and the Open Philanthropy Project, an advised fund of the Silicon Valley Community Foundation.

## Supplementary Information (SI)

**Table S1.** Nucleotide diversity and Tajima’s D for the six *An. gambiae* populations. The last row is all *An. gambiae* (2918) combined for comparison. Site choice was as described in the main text. The asterisk * denotes statistical significance, (p < 0.05). † Denotes an island population.

| Population | Nucleotide diversity ($\pi$) | Tajima's D |
| --- | --- | --- |
| Kayonjo | 0.01163 | -2.3069* |
| Katuuso | 0.01143 | -2.0219* |
| Kibbuye | 0.01169 | -2.2356* |
| Kiimi † | 0.01081 | -0.8223 |
| Kansambwe † | 0.01071 | -1.0827 |
| Bugiri † | 0.01103 | -0.6954 |
| All *An. gambiae* (Mainland & Island) | 0.01150 | -2.3073* |

**Table S2.** Nucleotide diversity and Tajima’s D for the three *An. arabiensis* populations The last row is all *An. arabiensis* (173) combined for comparison. Site choice was as described in the main text.

| Population | Nucleotide diversity ($\pi$) | Tajima's D |
| --- | --- | --- |
| Kayonjo | 0.00859 | -1.17475 |
| Katuuso | 0.00829 | -0.98993 |
| Kibbuye | 0.00847 | -1.73386 |
| All *An. arabiensis* | 0.00859 | -1.91156 |

## Notes

### Competing Interest Statement

The authors have declared no competing interest.

### Summary of Updates

Reviewers shared their comments which have been incoporated in the updated version

## References

1. Bhatt S, Weiss DJ, Cameron E, Bisanzio D, Mappin B, Dalrymple U, et al. The effect of malaria control on Plasmodium falciparum in Africa between 2000 and 2015. 2015;526:207–11.

2. Lehmann T, Dao A, Yaro AS, Adamou A, Kassogue Y, Diallo M, et al. Aestivation of the African Malaria Mosquito, Anopheles gambiae in the Sahel. American Journal of Tropical Medicine & Hygiene. 2010;83:601–6.

3. WHO. World Malaria Report. World Health Organization; 2024.

4. WHO. World Malaria Report. World Health Organization. 2023.

5. WHO. World Malaria Report. World Health Organization. 2022.

6. Hemingway J. The role of vector control in stopping the transmission of malaria: Threats and opportunities. Philosophical Transactions of the Royal Society B: Biological Sciences. Royal Society; 2014.

7. Mushtaq I, Sarwar MS, Chaudhry A, Shah SAH, Ahmad MM. Updates on traditional methods for combating malaria and emerging Wolbachia-based interventions. Front Cell Infect Microbiol. Frontiers Media SA; 2024.

8. Plowe C V. Malaria chemoprevention and drug resistance: a review of the literature and policy implications. Malar J. BioMed Central Ltd; 2022.

9. Thu AM, Phyo AP, Landier J, Parker DM, Nosten FH. Combating multidrug-resistant Plasmodium falciparum malaria. FEBS Journal. Blackwell Publishing Ltd; 2017. p. 2569–78.

10. Kaddumukasa MA, Wright J, Muleba M, Stevenson JC, Norris DE, Coetzee M. Genetic differentiation and population structure of Anopheles funestus from Uganda and the southern African countries of Malawi, Mozambique, Zambia and Zimbabwe. Parasit Vectors [Internet]. 2020;13:1–13. Available from: 10.1186/s13071-020-3962-1

11. Hammond A, Pollegioni P, Persampieri T, North A, Minuz R, Trusso A, et al. Gene-drive suppression of mosquito populations in large cages as a bridge between lab and field. Nat Commun. 2021;12.

12. Marshall JM, Akbari OS. Gene Drive Strategies for Population Replacement. Genetic Control of Malaria and Dengue. 2016;169–200.

13. Burt A. Heritable strategies for controlling insect vectors of disease. Philosophical Transactions of the Royal Society B: Biological Sciences. 2014;369.

14. Gantz VM, Jasinskiene N, Tatarenkova O, Fazekas A, Macias VM, Bier E, et al. Highly efficient Cas9-mediated gene drive for population modification of the malaria vector mosquito *Anopheles stephensi*. Proc Natl Acad Sci U S A. 2015;112:E6736–43.

15. Hammond MA, Galizi R. Gene drives to fight malaria: current state and future directions. Pathog Glob Health. 2017;111:412–23.

16. Makunin A, Korlević P, Park N, Goodwin S, Waterhouse RM, von Wyschetzki K, et al. A targeted amplicon sequencing panel to simultaneously identify mosquito species and Plasmodium presence across the entire *Anopheles* genus. Mol Ecol Resour. 2022;22:28–44.

17. Galizi R, Doyle LA, Menichelli M, Bernardini F, Deredec A, Burt A, et al. A synthetic sex ratio distortion system for the control of the human malaria mosquito. Nat Commun. 2014;5.

18. Gantz VM, Bier E. The mutagenic chain reaction: A method for converting heterozygous to homozygous mutations. Science (1979). 2015;348:442–4.

19. Hammond A, Galizi R, Kyrou K, Simoni A, Siniscalchi C, Katsanos D, et al. A CRISPR-Cas9 gene drive system targeting female reproduction in the malaria mosquito vector *Anopheles gambiae*. Nat Biotechnol. 2016;34:78–83.

20. Ng’habi, R., K., Knols, J., G., B., Lee, Y., Ferguson, M., H. and Lanzaro, C. G. Population genetic structure of Anopheles arabiensis and Anopheles gambiae in a malaria endemic region of southern Tanzania. Malar J. 2011;10.

21. Lanzaro GC, Tripet F. Gene flow among populations of Anopheles gambiae: a critical review. 2003;1–24. Available from: http://edepot.wur.nl/136907%5Cnpapers3://publication/uuid/F232B8BC-F74F-456E-A0DB-11E8198EE5B1

22. Lanzaro GC, Touré YT, Carnahan J, Zheng L, Dolo G, Traoré S, et al. Complexities in the genetic structure of *Anopheles gambiae* populations in west Africa as revealed by microsatellite DNA analysis. Proc Natl Acad Sci U S A. 1998;95:14260–5.

23. Burt A. Site-specific selfish genes as tools for the control and genetic engineering of natural populations. Proceedings of the Royal Society B: Biological Sciences. 2003;270:921– 8.

24. Naidoo K, Oliver S V. Gene drives: an alternative approach to malaria control? Gene Ther. Springer Nature; 2024.

25. Hancock PA, North A, Leach AW, Winskill P, Ghani AC, Godfray HCJ, et al. The potential of gene drives in malaria vector species to control malaria in African environments. Nat Commun [Internet]. 2024;15:8976. Available from: https://www.nature.com/articles/s41467-024-53065-z

26. Wiltshire MR, Bergey MC, Kayondo KJ, Birungi J, Mukwaya GL, Emrich JS, et al. Reduced-representation sequencing identifies small effective population sizes of *Anopheles gambiae* in the north-western Lake Victoria basin, Uganda. Malar J. 2018;17.

27. Kayondo, K., J., Mukwaya, G., L., Stump, A., Michel, P., A., Coulibaly, B., M., Besansky, J.,N. and Collins, H. F. Genetic structure of Anopheles gambiae populations on islands in northwestern Lake Victoria, Uganda. Malar J. 2005;4.

28. Kayondo, K., J., Mukwaya, G., L., Stump, A., Michel, P., A., Coulibaly, B., M., Besansky, J., N. and Collins, H. F. Genetic structure of Anopheles gambiae populations on islands in northwestern Lake Victoria, Uganda. Malar J. 2005;4.

29. Lukindu, M., Bergey, M., C., Wiltshire, M., R., Small, T., S., Bourke, P., B., Kayondo, K., J. and Besansky, J. N. Spatio-temporal genetic structure of Anopheles gambiae in the Northwestern Lake Victoria Basin, Uganda: implications for genetic control trials in malaria endemic regions. Parasit Vectors. 2018;11.

30. Wiltshire, M., R., Bergey, M., C., Kayondo, K., J., et al. Reduced-representation sequencing identifies small effective population sizes of Anopheles gambiae in the north-western Lake Victoria basin, Uganda. Malar J. 2018;17.

31. M. DJ, Cuamba N., Charlwood JD, F. CH, Townson H. Population structure in the malaria vector, Anopheles arabiensis Patton, in East Africa. Heredity (Edinb). 1999;83:408–17.

32. Della Torre A, Tu Z, Petrarca V. On the distribution and genetic differentiation of *Anopheles gambiae s.s.* molecular forms. Insect Biochem Mol Biol. 2005;35:755–69.

33. O’Loughlin SM, Magesa S, Mbogo C, Mosha F, Midega J, Lomas S, et al. Genomic analyses of three malaria vectors reveals extensive shared polymorphism but contrasting population histories. Mol Biol Evol. 2014;31:889–902.

34. Lukindu M, Bergey MC, Wiltshire MR, Small TS, Bourke PB, Kayondo KJ, et al. Spatio-temporal genetic structure of *Anopheles gambiae* in the Northwestern Lake Victoria Basin, Uganda: implications for genetic control trials in malaria endemic regions. Parasit Vectors. 2018;11.

35. Okech BA, Gouagna LC, Killeen GF, Knols BGJ, Kabiru EW, Beier JC, et al. Influence of sugar availability and indoor microclimate on survival of *Anopheles gambiae* (Diptera: Culicidae) under semifield conditions in western Kenya. J Med Entomol. 2003;40:657–63.

36. Ngowo HS, Kaindoa EW, Matthiopoulos J, Ferguson HM, Okumu FO. Variations in household microclimate affect outdoor-biting behaviour of malaria vectors [version 1; referees: 2 approved, 1 approved with reservations]. Wellcome Open Res. 2017;2:1–18.

37. Zhong D, Wang X, Xu T, Zhou G, Wang Y, Lee MC, et al. Effects of microclimate condition changes due to land use and land cover changes on the survivorship of malaria vectors in China-Myanmar border region. PLoS One. 2016;11.

38. Gillies MT, and, Coetzee M. A supplement to the *Anopheline* of Africa South of the Sahara (Afro-Tropical region). South African Institute for Medical Research. 1987;55.

39. Boddé M, Makunin A, Ayala D, Bouafou L, Diabaté A, Ekpo UF, et al. High-resolution species assignment of *Anopheles* mosquitoes using k-mer distances on targeted sequences. Elife. 2022;11:1–40.

40. Kingma DP, Welling M. Auto-Encoding Variational Bayes. 2013; Available from: http://arxiv.org/abs/1312.6114

41. Nagi SC, Lucas ER, Ashraf F, Mugoya T, Lukyamuzi E, Summers S, et al. Targeted genomic surveillance of insecticide resistance in African malaria vectors [Internet]. 2025. Available from: http://biorxiv.org/lookup/doi/10.1101/2025.02.14.637727

42. Ochoa A, Storey JD. FST and kinship for arbitrary population structures I: Generalized definitions [Internet]. 2016. Available from: http://biorxiv.org/lookup/doi/10.1101/083915

43. Ochoa A, Storey JD. Estimating FST and kinship for arbitrary population structures. PLoS Genet. 2021;17.

44. Pritchard KJ, Stephens M, Donnelly P. Inference of Population Structure Using Multilocus Genotype Data. Genetics. 2000;155:945–959.

45. Ochola A. SJD. popkinsuppl: supplement to popkin package [Internet]. 2022. Available from: https://github.com/OchoaLab/popkinsuppl.

46. Bhatia G, Patterson N, Sankararaman S, Price AL. Estimating and interpreting F ST: The impact of rare variants. Genome Res. 2013;23:1514–21.

47. Arnold B, Corbett-Detig RB, Hartl D, Bomblies K. RADseq underestimates diversity and introduces genealogical biases due to nonrandom haplotype sampling. Mol Ecol. 2013;22:3179–90.

48. Efron B, Tibshirani JR. An introduction to the Bootstrap. Cox DR, Hinkley D V, Reid N, Rubin DB, Silverman BW, editors. Regression Analysis with Applications B.G. Wetherill. Taylor & Francis Group; 1994.

49. Peakall R, Smouse PE. GenALEx 6.5: Genetic analysis in Excel. Population genetic software for teaching and research-an update. Bioinformatics. 2012;28:2537–9.

50. Diniz-Filho JAF, Soares TN, Lima JS, Dobrovolski R, Landeiro VL, Telles MP de C, et al. Mantel test in population genetics. Genet Mol Biol. 2013;36:475–85.

51. Patterson N, Price AL, Reich D. Population Structure and Eigenanalysis. PLoS Genet. 2006;2.

52. Team RC. R: A language and environment for statistical computing. R Foundation for statistical computing, Vienna, Austria. 2023.

53. Evanno G, Regnaut S, Goudet J. Detecting the number of clusters of individuals using the software STRUCTURE: A simulation study. Mol Ecol. 2005;14:2611–20.

54. Kopelman NM, Mayzel J, Jakobsson M, Rosenberg NA, Mayrose I. Clumpak: A program for identifying clustering modes and packaging population structure inferences across K. Mol Ecol Resour. 2015;15:1179–91.

55. Nei M, Miller JC. A simple method for estimating average number of nucleotide substitutions within and between populations from restriction data. Genetics. 1990;125:873–9.

56. Tajima F. Statistical Method for Testing the Neutral Mutation Hypothesis by DNA Polymorphism. Genetics Society of America. 1989;123:585–95.

57. Paradis E. Pegas: An R package for population genetics with an integrated-modular approach. Bioinformatics. 2010;26:419–20.

58. Waples, S., R. & Do C. LDNE: a program for estimating effective population size from data on linkage disequilibrium. Mol Ecol Resour. 2008;8:753–6.

59. Do C, Waples RS, Peel D, Macbeth GM, Tillett BJ, Ovenden JR. NeEstimator v2: Re-implementation of software for the estimation of contemporary effective population size (Ne) from genetic data. Mol Ecol Resour. 2014;14:209–14.

60. F. R. Genetic Differentiation and Estimation of Gene Flow from FStatistics Under Isolation by Distance. Genetics. 1997;34:399–405.

61. Evanno G, Regnaut S, Goudet J. Detecting the number of clusters of individuals using the software STRUCTURE: A simulation study. Mol Ecol. 2005;14:2611–20.

62. Service MW. Mosquito (Diptera: Culicidae) Dispersal-The Long and Short of It. J Med Entomol [Internet]. 1997;34:579–88. Available from: https://academic.oup.com/jme/article/34/6/579/2221644

63. Bataille A, Cunningham AA, Cruz M, Cedeño V, Goodman SJ. Adaptation, isolation by distance and human-mediated transport determine patterns of gene flow among populations of the disease vector Aedes taeniorhynchus in the Galapagos Islands. Infection, Genetics and Evolution. 2011;11:1996–2003.

64. Kayondo JK, Mukwaya LG, Stump A, Michel AP, Coulibaly MB, Besansky NJ, et al. Genetic structure of *Anopheles gambiae* populations on islands in northwestern Lake Victoria, Uganda. Malar J. 2005;4.

65. Atieli HE, Zhou G, Zhong D, Wang X, Lee MC, Yaro AS, et al. Wind-assisted high-altitude dispersal of mosquitoes and other insects in East Africa. J Med Entomol. 2023;60:698–707.

66. Sanogo ZL, Yaro AS, Dao A, Diallo M, Yossi O, Samaké D, et al. The effects of high-altitude windborne migration on survival, oviposition, and blood-feeding of the african malaria mosquito, Anopheles gambiae s.l. (Diptera: Culicidae). J Med Entomol. 2021;58:343–9.

67. Kalinowski ST. Do polymorphic loci require large sample sizes to estimate genetic distances? Heredity (Edinb). 2005;94:33–6.

68. Slatkin M. Gene Flow and the Geographic Structure of Natural Populations [Internet]. 1987. Available from: https://www.science.org

69. Charlesworth B. Measures of Divergence Between Populations and the Effect of Forces that Reduce Variability. Mol Biol Evol [Internet]. 1998;15:538–43. Available from: https://academic.oup.com/mbe/article/15/5/538/987854

70. Hudson RR, Slatkint M, Maddison WP. Estimation of Levels of Gene Flow From DNA Sequence Data. 1992.

71. Smaragdov MG, Kudinov AA, Uimari P. Assessing the genetic differentiation of holstein cattle herds in the Leningrad region using Fst statistics. Agricultural and Food Science. 2018;27:96–101.

72. Hartl DL, Clark AG. PRINCIPLES OF POPULATION GENETICS. 4th ed. Sinauer Associates; 2007.

73. Holsinger KE, Weir BS. Genetics in geographically structured populations: Defining, estimating and interpreting FST. Nat Rev Genet. 2009. p. 639–50.

74. Wright S. The genetical structure of populations. nnals of Eugenics. 1951;15:323–354.

75. North AR, Burt A, Godfray HCJ, North, R., A., Burt, A. and Godfray, J., H. C. Modelling the potential of genetic control of malaria mosquitoes at national scale. BMC Biol. 2019;17:1–12.

76. Unckless RL, Clark AG, Messer PW. Evolution of resistance against CRISPR/Cas9 gene drive. Genetics. 2017;205:827–41.

77. Das S, Máquina M, Phillips K, Cuamba N, Marrenjo D, Saúte F, et al. Fine-scale spatial distribution of deltamethrin resistance and population structure of *Anopheles funestus* and *Anopheles arabiensis* populations in Southern Mozambique. Malar J. 2023;22.

78. Fumagalli M. Assessing the effect of sequencing depth and sample size in population genetics inferences. PLoS One. 2013;8.

79. Bi K, Linderoth T, Singhal S, Vanderpool D, Patton JL, Nielsen R, et al. Temporal genomic contrasts reveal rapid evolutionary responses in an alpine mammal during recent climate change. PLoS Genet. 2019;15.

80. Machado HE, Bergland AO, Taylor R, Tilk S, Behrman E, Dyer K, et al. Broad geographic sampling reveals the shared basis and environmental correlates of seasonal adaptation in drosophila. Elife. 2021;10.

81. Sandoval-Castellanos E. Testing temporal changes in allele frequencies: A simulation approach. Genet Res (Camb). 2010;92:309–20.

82. Diniz-Filho JAF, Soares TN, Lima JS, Dobrovolski R, Landeiro VL, Telles MP de C, et al. Mantel test in population genetics. Genet Mol Biol. 2013;36:475–85.

83. The Anopheles gambiae 1000 Genomes consortium. Genetic diversity of the African malaria vector Anopheles gambiae. Nature. 2017;552:96–100.

84. Guillot G. Inference of structure in subdivided populations at low levels of genetic differentiation - The correlated allele frequencies model revisited. Bioinformatics. 2008;24:2222–8.

85. Weir BS, Cockerham CC. Estimating F-statistics for the analysis of population structure. Evolution (N Y). 1984;38:1358–70.

86. Jombart T, Devillard S, Balloux F. Discriminant analysis of principal components: A new method for the analysis of genetically structured populations. BMC Genet. 2010;11.

87. Nielsen R. Molecular signatures of natural selection. Annu Rev Genet. 2005. p. 197–218.

88. Kreitman M. Methods to detect selection in populations with applications to the human. Annu Rev Genomics Hum Genet. 2000;1:539–59.

89. Tajima F. Statistical Method for Testing the Neutral Mutation Hypothesis by DNA Polymorphism. 1989.

90. O’Loughlin SM, Magesa S, Mbogo C, Mosha F, Midega J, Lomas S, et al. Genomic analyses of three malaria vectors reveals extensive shared polymorphism but contrasting population histories. Mol Biol Evol. 2014;31:889–902.

91. Nei M, Li WH. Mathematical model for studying genetic variation in terms of restriction endonucleases. Proc Natl Acad Sci U S A. 1979;76:5269–73.

92. Parimittr P, Chareonviriyaphap T, Bangs MJ, Arunyawat U. Genetic variation of Aedes aegypti mosquitoes across Thailand based on nuclear DNA sequences. Agriculture and Natural Resources. 2018;52:596–602.

93. Dharmarathne HAKM, Weerasena OVDSJ, Perera KLNS, Galhena G. Genetic characterization of Aedes aegypti (Diptera: Culicidae) in Sri Lanka based on COI gene. J Vector Borne Dis. 2020;57:153–60.

94. Besansky NJ, Krzywinski J, Lehmann T, Simard F, Kernt M, Mukabayire O, et al. Semipermeable species boundaries between *Anopheles gambiae* and *Anopheles arabiensis*: Evidence from multilocus DNA sequence variation. Proc Natl Acad Sci U S A. 2003;100:10818–23.

95. Obbard DJ, Linton YM, Jiggins FM, Yan G, Little TJ. Population genetics of Plasmodium resistance genes in *Anopheles gambiae*: No evidence for strong selection. Mol Ecol. 2007;16:3497–510.

96. Campos M, Hanemaaijer M, Gripkey H, Collier TC, Lee Y, Cornel AJ, et al. The origin of island populations of the African malaria mosquito, Anopheles coluzzii. Commun Biol. 2021;4.

97. Bergey CM, Lukindu M, Wiltshire RM, Fontaine MC, Kayondo JK, Besansky NJ. Assessing connectivity despite high diversity in island populations of a malaria mosquito. Evol Appl. 2020;13:417–31.

98. Lanzaro GC, Campos M, Crepeau M, Cornel A, Estrada A, Gripkey H, et al. Selection of sites for field trials of genetically engineered mosquitoes with gene drive. Evol Appl. 2021;14:2147–61.

99. Ellegren H, Galtier N. Determinants of genetic diversity. Nat Rev Genet. 2016;17:422–33.

100. Fischer MC, Rellstab C, Leuzinger M, Roumet M, Gugerli F, Shimizu KK, et al. Estimating genomic diversity and population differentiation - an empirical comparison of microsatellite and SNP variation in Arabidopsis halleri. BMC Genomics. 2017;18:1–15.

101. O’Loughlin SM, Magesa SM, Mbogo C, Mosha F, Midega J, Burt A. Genomic signatures of population decline in the malaria mosquito *Anopheles gambiae*. Malar J. 2016;15:1–10.

102. Lobo NF, Sangaré DM, Regier AA, Reidenbach KR, Bretz DA, Sharakhova M V., et al. Breakpoint structure of the *Anopheles gambiae* 2Rb chromosomal inversion. Malar J. 2010;9.

103. White BJ, Cheng C, Sangaré D, Lobo NF, Collins FH, Besansky NJ. The population genomics of trans-specific inversion polymorphisms in *Anopheles gambiae*. Genetics. 2009;183:275–88.

104. Petrarca V, Nugud AD, Elkarim Ahmed MA, Haridi AM, Di Deco MA, Coluzzi M. Cytogenetics of the Anopheles gambiae complex in Sudan, with special reference to An. arabiensis: Relationships with East and West African populations. Med Vet Entomol. 2000;14:149–64.

105. Peter Andolfatto, Frantz Depaulis AN. Inversion polymorhisms and nucleotide variability in Drosophila. Genetics Research Cambridge. 2001;77:1–8.

106. Hare MP, Nunney L, Schwartz MK, Ruzzante DE, Burford M, Waples RS, et al. Understanding and Estimating Effective Population Size for Practical Application in Marine Species Management. Conservation Biology. 2010. p. 438–49.

107. Waples RS, Do C. Linkage disequilibrium estimates of contemporary Ne using highly variable genetic markers: A largely untapped resource for applied conservation and evolution. Evol Appl. 2009;3:244–62.

108. Wang J. Estimation of effective population sizes from data on genetic markers. Philosophical Transactions of the Royal Society B: Biological Sciences. Royal Society; 2005. p. 1395–409.

109. Rousset F. Genetic differentiation and estimation of gene flow from F-statistics under isolation by distance. Genetics. 1997;145:1219–28.

110. Neafsey DE, Waterhouse RM. Highly evolvable malaria vectors: The genomes of 16 Anopheles mosquitoes. 2015 [cited 2024 Aug 12]; Available from: http://dx.doi.

111. Hoffmann AA, Willi Y. Detecting genetic responses to environmental change. Nat Rev Genet. 2008. p. 421–32.

112. Weetman D, Wilding CS, Neafsey DE, Müller P, Ochomo E, Isaacs AT, et al. Candidate-gene based GWAS identifies reproducible DNA markers for metabolic pyrethroid resistance from standing genetic variation in East African *Anopheles gambiae*. Sci Rep. 2018;8.

113. Athrey G, Hodges TK, Reddy MR, Overgaard HJ, Matias A, Ridl FC, et al. The Effective Population Size of Malaria Mosquitoes: Large Impact of Vector Control. PLoS Genet. 2012;8.

114. Charlesworth B. Effective population size and patterns of molecular evolution and variation. Nature Reviews|Genetics. 2009;10:195–205.

115. Yaro AS, Traoré AI, Huestis DL, Adamou A, Timbiné S, Kassogué Y, et al. Dry season reproductive depression of *Anopheles gambiae* in the Sahel. J Insect Physiol [Internet]. 2012;58:1050–9. Available from: 10.1016/j.jinsphys.2012.04.002

116. White MT, Griffin JT, Churcher TS, Ferguson NM, Basáñez MG, Ghani AC. Modelling the impact of vector control interventions on *Anopheles gambiae* population dynamics. Parasit Vectors. 2011;4.

117. Kamdem C, Fouet C, Gamez S, White BJ. Pollutants and Insecticides Drive Local Adaptation in African Malaria Mosquitoes. Mol Biol Evol. 2017;34:1261–75.

118. Waples RS. Life-history traits and effective population size in species with overlapping generations revisited: The importance of adult mortality. Heredity (Edinb). 2016;117:241–50.

119. Mwima R, Hui TYJ, Kayondo JK, Burt A. The population genetics of partial diapause, with applications to the aestivating malaria mosquito *Anopheles coluzzii*. Mol Ecol Resour. 2024;24.

120. Mwima R, Hui TYJ, Nanteza A, Burt A, Kayondo JK. Potential persistence mechanisms of the major *Anopheles gambiae* species complex malaria vectors in sub-Saharan Africa: a narrative review. Malar J [Internet]. 2023 [cited 2024 Jun 24];22:1–12. Available from: https://malariajournal.biomedcentral.com/articles/10.1186/s12936-023-04775-0

